# Nitrous oxide modulates cortical activity, wake–sleep oscillations, and produces antidepressant-like effects in mice

**DOI:** 10.1101/2025.02.13.638096

**Authors:** Samuel Kohtala, Puja K Parekh, Iman Baramaki, Piia Kohtala, Liisa Ilkka, Hanna Antila, Francis S. Lee, Conor Liston

**Affiliations:** Department of Psychiatry, Weill Cornell Medicine, New York, NY, USA; Drug Research Program, Division of Pharmacology and Pharmacotherapy, Faculty of Pharmacy, University of Helsinki, Finland; Neuroscience Center and SleepWell Research Program, Faculty of Medicine, University of Helsinki, Finland

**Keywords:** rapid-acting antidepressant, nitrous oxide, N2O, calcium imaging, EEG, sleep, chronic corticosterone

## Abstract

Emerging evidence suggests that nitrous oxide (N_2_O), a gaseous NMDA receptor antagonist and dissociative anesthetic, exerts rapid antidepressant effects akin to subanesthetic ketamine. However, its cellular, molecular, and behavioral effects remain poorly understood. Using in vivo two-photon imaging through cortical microprisms, we demonstrate that 50% N_2_O/O_2_ rapidly increases neuronal calcium activity in the mouse medial prefrontal cortex (mPFC). This was corroborated by elevated c-Fos expression at both protein and mRNA levels in mPFC lysates. Cortical EEG recordings revealed that N_2_O increased subsequent wake-associated gamma oscillations and enhanced slow-wave activity during sleep, suggestive of cortical activation and synaptic potentiation. In a chronic corticosterone stress model, N_2_O elicited antidepressant-like behavioral effects in several, though not all, domains. Together, these findings indicate that a single treatment with N_2_O rapidly enhances cortical activity, modulates sleep and wake EEG oscillations, and produces antidepressant-like effects, paralleling key actions associated with subanesthetic ketamine.

## Introduction

Nitrous oxide (N_2_O), a gaseous dissociative anesthetic, has recently emerged as a potential rapid-acting treatment for depression (Desmidt et al., 2023; Liu et al., 2023; Nagele et al., 2021, Nagele et al., 2015; Shao et al., 2023). N_2_O acts as an antagonist of N-methyl-D-aspartate receptors (NMDARs), which is the primary pharmacological mechanism of action of the rapid-acting antidepressant drug ketamine (Kohtala, 2021). Unlike ketamine, N_2_O does not undergo systemic metabolism and thereby has limited potential for drug interactions, while showing consistent rapid pharmacokinetics (Quach et al., 2022). Moreover, the pharmacological effects of N_2_O become evident almost instantly upon inhalation and recover rapidly upon cessation of gas flow, which has advantages for clinical treatment (Kalmoe et al., 2020). However, the cellular and molecular mechanisms underlying the effects of N_2_O remain to be thoroughly characterized.

The similarities in the pharmacological mode of action of ketamine and N_2_O raises the question whether their subsequent cellular and molecular effects also have shared features (Kohtala and Rantamäki, 2021). Subanesthetic doses of ketamine are thought to preferentially inhibit NMDARs in gamma-aminobutyric acidergic (GABAergic) interneurons, resulting in the disinhibition of excitatory pyramidal neurons and increase in glutamate release and bursting (Ali et al., 2020; Homayoun and Moghaddam, 2007; Miller et al., 2016; Moghaddam et al., 1997; Widman and McMahon, 2018). Ketamine has been shown to acutely increase medial prefrontal cortex (mPFC) calcium activity, electroencephalography (EEG) gamma power, and the expression of the activity-dependent immediate-early gene (IEG) c-Fos (Davoudian et al., 2023; Hare et al., 2020; Kohtala et al., 2019c; Li et al., 2010, 2010; Salort et al., 2019). Notably, recent studies indicate that both N_2_O and ketamine persistently enhance excitatory transmission in CA1 hippocampal slices (Izumi et al., 2022) and upregulate molecular markers of neuronal activity in the mouse mPFC (Kohtala et al., 2019c; Rozov et al., 2024), which suggests potential converging mechanisms underlying their effects.

In the present study, we investigated the effects of a single N_2_O exposure, at a clinically used concentration (50% N_2_O/O_2_), on neuronal activity by utilizing *in vivo* two-photon calcium imaging, quantitative cortical EEG recordings, and a variety of molecular techniques. Furthermore, we utilized the chronic corticosterone model to test whether N_2_O treatment produces antidepressant-like effects across a battery of behavioral tests. Our findings indicate that N_2_O treatment rapidly increases calcium activity and molecular markers indicative of cortical activation, influences EEG features associated with synaptic plasticity, and produces rapid antidepressant-like effects in a model of chronic stress. The findings provide insights into the effects of N_2_O on neuronal function and highlight the potential of N_2_O as an experimental tool to investigate mechanisms underlying rapid antidepressant effects in general.

## Materials and methods

### Animals

Adult male C57BL/6J mice (aged 9-15 weeks) were used for all experiments. Animals were maintained in the animal facility under standard conditions (21 °C, 12-h light-dark cycle, lights on 06:00 and off 18:00) with free access to food and water (except during sucrose preference test). Experiments were conducted during the light (active) period, between 09:00-16:00. The experiments were carried out according to the guidelines of the Society for Neuroscience and were approved by the Weill Cornell Institutional Animal Care and Use Committee, or the European Communities Council Directive of 22 September 2010 (Directive 2010/63/EU) and approved by the County Administrative Board of Southern Finland (ESAVI/5844/2019).

### Stereotactic surgery and viral vectors

Animals were anesthetized using isoflurane anesthesia (induction 4%, maintenance 1-2%) and treated with metacam (1 mg/kg, i.p.) as prophylactic analgesic. Animals were placed in a stereotactic frame, eyes were moisturized to prevent corneal drying and microwavable heating pad was used to maintain body temperature. Scalp fur was trimmed, and bupivacaine was administered (0,05 ml, 5 mg/ml) at the site of incision, and the skull surface was exposed. The center of the target site for injection was determined using stereotaxic coordinates from bregma (+1,7AP, 0,35ML, −1,35DV), corresponding to the cingulate cortex. AAV1-hSyn-GCaMP6s (a gift from Douglas Kim & GENIE Project (Addgene plasmid # 100843; http://n2t.net/addgene:100843; RRID:Addgene_100843) was injected into the mPFC for two-photon imaging. The virus was injected at a rate of 100 nl/min using a Nanofil syringe (World Precision Instruments) with a 33 G beveled needle (World Precision Instruments) and injector (World Precision Instruments) at a total volume of 500 nl. The needle kept in place for two minutes prior to its slow removal. For microprism implantation, a craniotomy was made close to the midline, and right-angle borosilicate glass prism (1,5×1,5×3mm) was implanted in contralateral to the viral injection site. The microprism was sealed to the skull using veterinary adhesive (Vetbond, Fisher Scientific), followed by a layer of dental cement (C&B Metabond, Parkell Inc). Finally, a custom-made circular titanium headplate was attached to the skull using Metabond. Upon completing the experiment, the mice were deeply anesthetized and underwent transcardial perfusion with phosphate-buffered saline (PBS), followed by 4% paraformaldehyde in PBS for histological analysis.

### Pharmacological treatments

Medical grade N_2_O (50% N_2_O/O_2_ mix, Airgas inc., USA, and Woikoski Oy, Finland) was used in all experiments. After habituation to the experimental conditions, the gas was administered either into acrylic chambers (for biochemical and behavioral experiments), into a modified home cage (EEG recordings), or directly using a nosecone (head fixed two-photon imaging). Medical grade oxygen and air were used as sham treatments.

### Chronic stress and behavioral tests

#### Chronic corticosterone (CORT)

Corticosterone (Sigma) was dissolved in 99,5% ethanol and diluted with animal facility-provided drinking water to a final concentration of 0,1 mg/ml of corticosterone and 1% ethanol (CORT). Mice were exposed to CORT in place of regular drinking water for 21 days. A control group received regular drinking water without CORT. Fresh solutions were changed twice a week.

#### Behavioral tests

Behavioral tests were performed at the following time point after treatment: open-field test (1h), sucrose preference (12h), tail suspension (24h), female urine sniffing (48h), coat score (48h). Mice were put into the test room at least 30 minutes before testing for habituation. All tests were performed between 9:00 - 16:00. in a quiet room dedicated for behavioral experiments. The animals were tested in a pseudorandomized manner so that animals belonging to different experimental groups were equally divided across the testing period. Experimental arenas were cleaned between subjects using 70% ethanol. Analyses were performed blind to the experimental conditions.

#### Open field test

The apparatus consisted of a light square box (45 × 45 × 45 cm) and an overhead camera. Mice were placed into the center of the box under dim light (200 lx) and were allowed to explore the arena for 10 minutes. A video-computerized tracking system (EthoVision) was used to record the distance traveled and center entries.

#### Tail suspension test (TST)

The mouse tail was inserted through a cut 1 ml pipette tip to prevent tail climbing and paper tape was used to attach the mice hanging upside down. The immobility time of each mouse was recorded over a 6 min period under dim light (200 lx). Movement was determined as major motion of the limbs or body, excluding minor movements of single limbs, or swinging motion resulting from previous momentum.

#### Female urine sniffing test (FUST)

The mice were first habituated to a cotton swab by attaching it to their home cage, pointing at the middle of the cage for 30 min. Mice were then habituated to the test by placing them into a fresh cage with the cotton swab moistened with water, and were allowed to explore for 5 min. For testing, the cotton swab was moistened with 50 ul of fresh urine collected from female mice, and the behavior was recorded for 5 min. Time spent sniffing was determined as the total duration the nose of the mouse was in close proximity to the tip of the swab. Biting of the tip, as well as climbing and other movements not directed towards the tip were excluded.

#### Sucrose preference

Mice were habituated to two water bottles in their home cage for 3 days before testing. On the test day, mice were restricted from access to water for 8 h prior to testing and temporarily single-housed in a cage containing standard water or 3% sucrose dissolved in water. Mice were allowed free access to the two water bottles for 12 hours throughout the active period. Water and sucrose water consumption was determined by weighing the bottles before and after the testing session, and sucrose preference was calculated as the fraction (%) of sucrose water consumption divided by total water consumption.

#### Coat state

Coat state was visually assessed by two independent evaluators blinded to the condition. The condition of their head, neck, dorsal coat, tail, forelimbs, hindlimbs, ventral coat and genital region was rated as 0 (normal) or 1 (abnormal). Total scores of the two evaluators were averaged for the final coat state score for each mouse. Abnormal score was assigned when the inspected region was visibly dirty, uneven, or rough.

#### 4-Choice task

The 4-choice odor task was performed essentially as previously described (Lu et al., 2021). A four chamber white acrylic arena 12” x 12” x 9” (L x W x H) with 4 quadrants partially divided by internal transparent walls was used. White ceramic ramekins (diameter = 2.88”, depth = 1.75”) were used to present the food reward and the odor stimuli. Used odors were essential oils (rosemary, thyme, clove, nutmeg, or cinnamon). Food rewards were Honey Nut Cheerios cut into pieces weighing approximately 10 mg each. Pine shavings were used to hide the food rewards in the ramekins. Between trials, the mouse was confined to a removable transparent acrylic cylinder (diameter = 6”) in the center of the arena.

Mice were placed on a food restricted diet for 5 days before the testing day to achieve a weight reduction of ~85% of initial weight. Mice were handled daily for habituation, their weight was monitored, and food allotment was served in the same empty ramekins used for the testing. On day 3 (habituation), the mouse was allowed to explore the arena and eat food rewards from ramekins (without pine shavings) in each quadrant for 10 min, repeated six times. On day 4 (shaping), only one ramekin with a reward was placed in one quadrant, and it was rotated between trials to reward all quadrants equally. First 4 trials the cereal was not covered, followed by 4 trials of dusting of shavings to 4 trials of quarter full, 4 trials of half full, and 12 trials of full coverage of the ramekin. On day 5 (testing), each mouse was tested for discrimination followed by a reversal session. Each quadrant had one ramekin filled with shavings and scented by a drop of essential oil on a piece of filter paper taped to the side of the ramekin. During the discrimination, one ramekin contained the reward and the mouse discriminated between the odors and learned to associate a certain odor with the ramekin containing the buried reward. The ramekins were pseudorandomized across trials. In each trial, the mouse was allowed to freely explore the quadrants until it started to dig a ramekin. If the choice was correct, the trial was terminated after the mouse had eaten the food reward, and if the choice was incorrect, the trial was terminated following digging. If the mouse did not commit to any ramekin within 3 min, the trial was terminated and considered an omission. After each trial, the mouse was returned to the center cylinder, ramekins were rearranged, and the food reward restored. The session criterion was met when the mouse correctly finished 8 out of 10 consecutive trials. Discrimination was directly followed by reversal, where the thyme odor was swapped for a novel odor (cinnamon), and the reward containing odor was switched from rosemary to clove. Reversal was carried out in a similar manner with the same session criterion.

### Dissection and processing of brain samples

Animals were euthanized at indicated times after the treatments by rapid cervical dislocation followed by decapitation. No anesthesia was used due to its potential confounding effects on the molecular analyses (Alitalo et al., 2023; Kohtala et al., 2016). Bilateral medial prefrontal cortex (including prelimbic and infralimbic cortices) was rapidly dissected on a cooled dish and stored at −80 °C.

For western blotting, the brain samples were homogenized in RIPA lysis buffer (Thermo Scientific) with added complete protease inhibitor mix (Roche) and PhosStop (Roche). After a 15 min incubation on ice, samples were centrifuged (16,000×g, 15 min, + 4 °C) and the resulting supernatant collected for further analysis. For the preparation of crude synaptosomes, brain samples were homogenized in 10% (w/v) ice-cold buffer containing 0.32 M sucrose, 20 mM HEPES pH 7.4, 1 mM EDTA, with added with protease and phosphatase inhibitors. Briefly, after centrifugation of the homogenate at 2800 rpm for 10 min at 4 °C, the supernatant was transferred to a new tube and centrifuged at 12 000 rpm for 10 min. The supernatant (cytosolic fraction) was removed and the resultant pellet, designated the crude synaptosomal fraction, was resuspended in RIPA lysis buffer with protease and phosphatase inhibitors. Sample protein concentrations were measured using Pierce BCA Protein assay kit (Thermo Scientific) or Bio-Rad DC protein assay (Bio-Rad Laboratories, Hercules, CA).

### Western blotting

Proteins (30–40 μg) were separated with SDS-PAGE under reducing and denaturing conditions and blotted to a polyvinylidene difluoride (PVDF) membrane using standard protocols. After blocking, the membranes were incubated with the following primary antibodies overnight: anti-GAPDH (#2118; 1:10 000, CST), anti-β-actin (#4967: 1:10 000, CST), anti-BDNF (ab108319; 1:1000, Abcam), anti-GluN1 (#5704; 1:1000, CST), anti-c-Fos (#4384, 1:1000, CST). Following primary antibody, the membranes were washed with TBS/0.1% Tween (TBST) and incubated with horseradish peroxidase conjugated secondary antibodies (1:10000, 1 h at room temperature; Bio-Rad). After subsequent washes, secondary antibodies were visualized using enhanced chemiluminescence (ECL or ECL Plus, Thermo Scientific) for detection by Biorad ChemiDoc MP camera (Bio-Rad Laboratories). ImageJ was used for quantifications of chemiluminescence.

### Quantitative RT-PCR

For samples collected from stress experiments, RNA was extracted using the RNAqueous-Micro Total RNA Isolation Kit (Invitrogen), and RNA concentration and purity was assessed with Nanodrop 2000 Spectrophotometer (Millipore) and diluted with nuclease free water to a concentration of 12 ng/ul. Samples with an RNA concentration of >100 ng/ul, and 260/280 nm ratio of >2.0 were processed further. cDNA was synthesized using the High-Capacity RNA-to-cDNA kit (Applied Biosystems) according to the manufacturer’s instructions. PCR was carried out using SYBR Green PCR Master Mix (Power) and diluted primer pairs (Table S1) that were separately validated. Each plate included both RT and NTC controls, and samples in triplicates. Samples were normalized to Gapdh transcript levels. Data was analyzed with QuantStudio 6 Flex Real-Time PCR System v1.3 (Applied Biosystems) and relative quantifications with the delta-delta Ct method. For samples collected after acute N_2_O exposure in naïve animals, qPCR was performed as previously described (Kohtala et al., 2019c).

### In vivo 2-photon calcium imaging

All in vivo images were acquired by using a 2P laser-scanning microscope (Olympus RS) with a Mai Tai DeepSee laser tuned to 920 nm and a 10x water-immersion objective (Olympus XLPLN10XSVMP). For calcium imaging experiments, time-lapse images were acquired using galvo scanning at ~1 frame per second in awake head-fixed mice.

Data was analyzed in MATLAB using the EzCalcium toolbox (Cantu et al., 2020). Imaging data was motion corrected using the integrated NoRMCorre algorithm (Pnevmatikakis and Giovannucci, 2017) and regions of interest (ROIs) were detected using CaImAn (Giovannucci et al., 2019). The following settings were used: initialization (Greedy), search method (Ellipse), merge threshold (0.9), fudge factor (0.95), spatial downsampling (1), temporal downsampling (1), temporal iterations (2). A MATLAB file was generated containing the extracted fluorescence (dF/F) for each ROI. Detected ROIs were refined using the ROI Refinement module, where baseline location was set to 15% and baseline time window was set as 10 seconds worth of frames. Exclusion criteria were set to roundness (0.5), oblongness (2), borderline (10), borderline allowance (1). Borderline ROIs were individually evaluated by the experimenter and ROIs displaying unusual shapes were excluded.

### Electroencephalography recordings

For implantation of the recording electrodes, mice were deeply anesthetized with isoflurane (4% induction, 1.5-2% maintenance), injected with 5mg/kg carprofen (s.c.) and local anesthetic lidocaine (s.c.) and attached to a stereotaxic frame. The body temperature was maintained with heating pad during the surgery. Three stainless steel screw electrodes connected via coated stainless-steel wires to a 6-pin connector, were attached to drilled holes in three locations (frontal, parietal and reference) in the skull. Two EMG wires were inserted in the neck musculature. Dental cement was used to secure the implant in place. The mice were allowed to recover for 2 weeks after the surgery, after which they were habituated to their individual recording cages and to the sleep recording setup for 3-5 days. During and after 50% N2O/O2 or sham (air) exposure, EEG and EMG signals were acquired with RHD2132 headstage (Intan Technologies) linked to Open Ephys acquisition board (Open Ephys).

For sleep analysis and to determine the brain state, we first computed the EEG and EMG spectrograms using sliding, half-overlapping 5 s windows, resulting in a time resolution of 2.5 s. Within each 5 s window, we estimated the power spectral density (PSD) using Welch’s method with a Hann window applied to sliding, half-overlapping 2 s intervals. Detailed description of the classifier used to detect the brain state and manual scoring GUI can be found in (Antila et al., 2022). All the annotations were manually confirmed or corrected by experienced annotator.

The power spectral density (PSD) of the EEG was computed using Welch’s method with Hann window for consecutive 3 s, half overlapping intervals. To calculate the power within a given frequency band, we approximated the corresponding area under the spectral density curve using a midpoint Riemann sum. Detailed description of the spindle detection algorithm can be found in (Antila et al., 2022).

### Statistics

All statistical analyses were performed with Graphpad Prism software (Versions 8 and 9; La Jolla, CA, USA) or with Pingouin statistical library for Python. A p ≤ 0.05 was considered statistically significant. Detailed descriptions of statistical tests for each experiment are shown in **Table S1**.

## Results

### Nitrous oxide increases calcium activity in the medial prefrontal cortex

Previous reports have demonstrated that subanesthetic ketamine acutely increases neuronal calcium activity in the prefrontal cortex (PFC) (Ali et al., 2020; Hare et al., 2020). To investigate whether nitrous oxide (N_2_O) exerts similar effects, we performed two-photon calcium imaging of GCaMP6s-expressing (Chen et al., 2013) layer 2/3 neurons in the medial PFC (mPFC) through a contralaterally implanted optical microprism. Mice were head-fixed with a nose cone providing 50% N_2_O/O_2_, and calcium activity was monitored over three sequential 7-minute phases (baseline, N_2_O, and recovery; **Figure 1A**).

**Figure 1.**
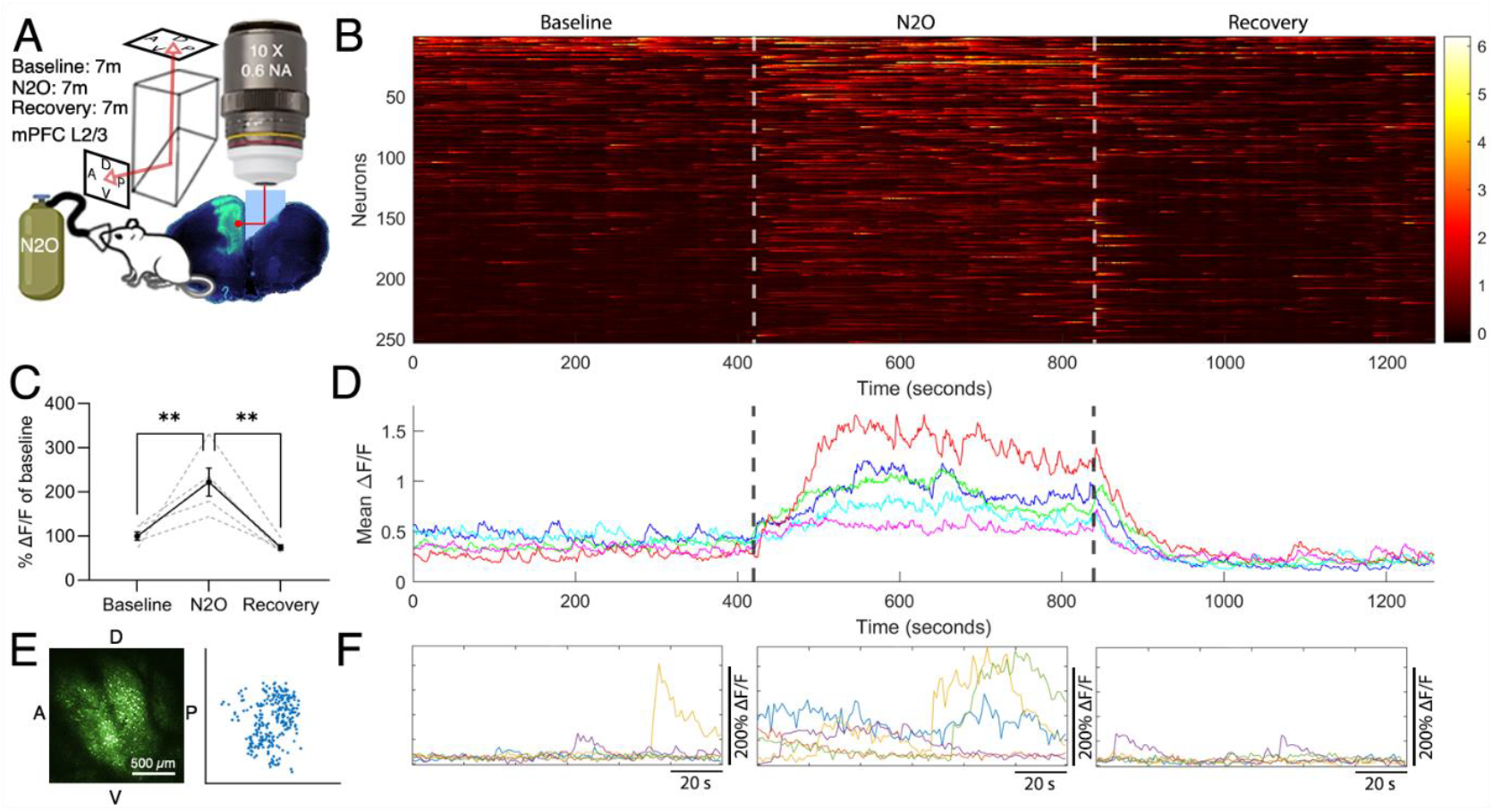
Effects of N_2_O on mPFC calcium dynamics. **A)** Imaging was directed to mPFC GCaMP6s expressing layer 2/3 neurons utilizing a contralateral microprism. Baseline calcium activity was determined, followed by N2O (50% N_2_O/O_2_) treatment, and recovery, each corresponding to 7 min of imaging for a total of 21 min. **B)** Representative heatmap of neuronal calcium traces in one subject. **C)** Mean calcium activity corresponding to baseline, N2O, and recovery periods across all subjects (n = 5), normalized to baseline. **D)** Averaged calcium traces from individual subjects. **E)** Representative cell map displaying neurons identified for one subject. **F)** Representative calcium traces from 5 randomly selected neurons of one subject during the three phases. **:p<0.01.

Regions of interest corresponding to individual neurons were automatically detected from AAV-hSyn-GCaMP6s signals in five subjects. Compared to baseline, N_2_O exposure elicited a rapid and statistically significant increase in mean neuronal calcium activity, which then subsided during the recovery period (**Figure 1B–F**). Importantly, control experiments involving sham treatments (either O_2_ or room air) failed to produce any notable increases in calcium activity (**Figure S1**), indicating that the observed increase in signal can be attributed to pharmacological action of N_2_O. Additional experiment with prolonged N_2_O exposure indicated that the enhancement of calcium activity was sustained over a longer exposure, only gradually diminishing over time (**Figure S2**). Collectively, these findings demonstrate that a brief exposure to a clinically relevant concentration of N_2_O rapidly increases neuronal calcium activity in the mouse mPFC.

### Nitrous oxide increases molecular markers of neuronal activity

Given that N_2_O robustly elevated calcium activity in the mPFC following a brief exposure, we next examined whether this effect corresponds to increased expression of immediate early genes (IEGs) associated with neuronal activation. Mice received 1 hour of either 50% N_2_O/O_2_ or sham (air) treatment, after which mPFC tissue was collected (**Figure 2**). We focused on the IEGs *Cfos* and *Arc*, which are rapidly upregulated following strong synaptic stimulation (Sagar et al., 1988; Steward and Worley, 2001). Western blot analysis revealed that c-Fos protein levels were significantly upregulated in the N_2_O group compared to sham (**Figure 2A**). In line with these findings, qPCR analysis showed a significant increase in *Cfos* mRNA, as well as a trend toward increased *Arc* mRNA expression (**Figure 2B**). These data suggest that N_2_O triggers robust neuronal activation in the mPFC that parallels ketamine’s reported effect on IEGs (Choi et al., 2015; Davoudian et al., 2023).

**Figure 2.**
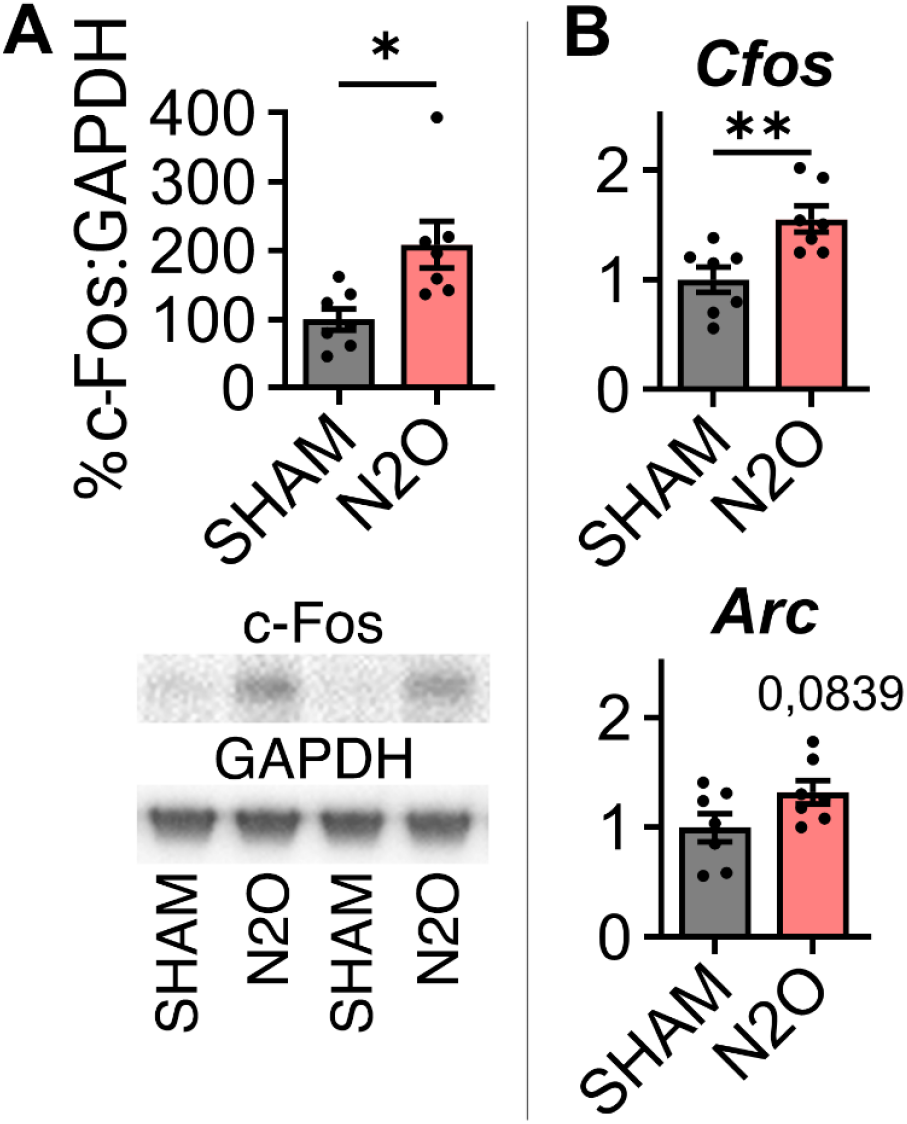
Effects of sham and N_2_O treatment on molecular markers of neuronal activation in the medial prefrontal cortex following 1 h treatment. **A)** Levels of c-Fos protein in the mPFC. **B)** Expression of *Cfos* and *Arc* mRNA. *:p<0.05, **:p<0.01.

### Nitrous oxide influences wake and sleep electroencephalography activity

Subanesthetic ketamine has previously been linked to alterations in cortical EEG oscillations, particularly enhanced gamma power, and to changes in sleep architecture that may contribute to its antidepressant properties (Gilbert and Zarate, 2020; Kohtala et al., 2020). To study whether N_2_O has similar effects in mice, we used EEG recordings to monitor cortical brain oscillations acutely during a 1h exposure to 50% N_2_O/O_2_ or sham, and for 2 h following treatment (**Figure 3A**). The same animals underwent both N_2_O and sham treatments in random order, with a minimum washout period of 1 week between treatments.

**Figure 3.**
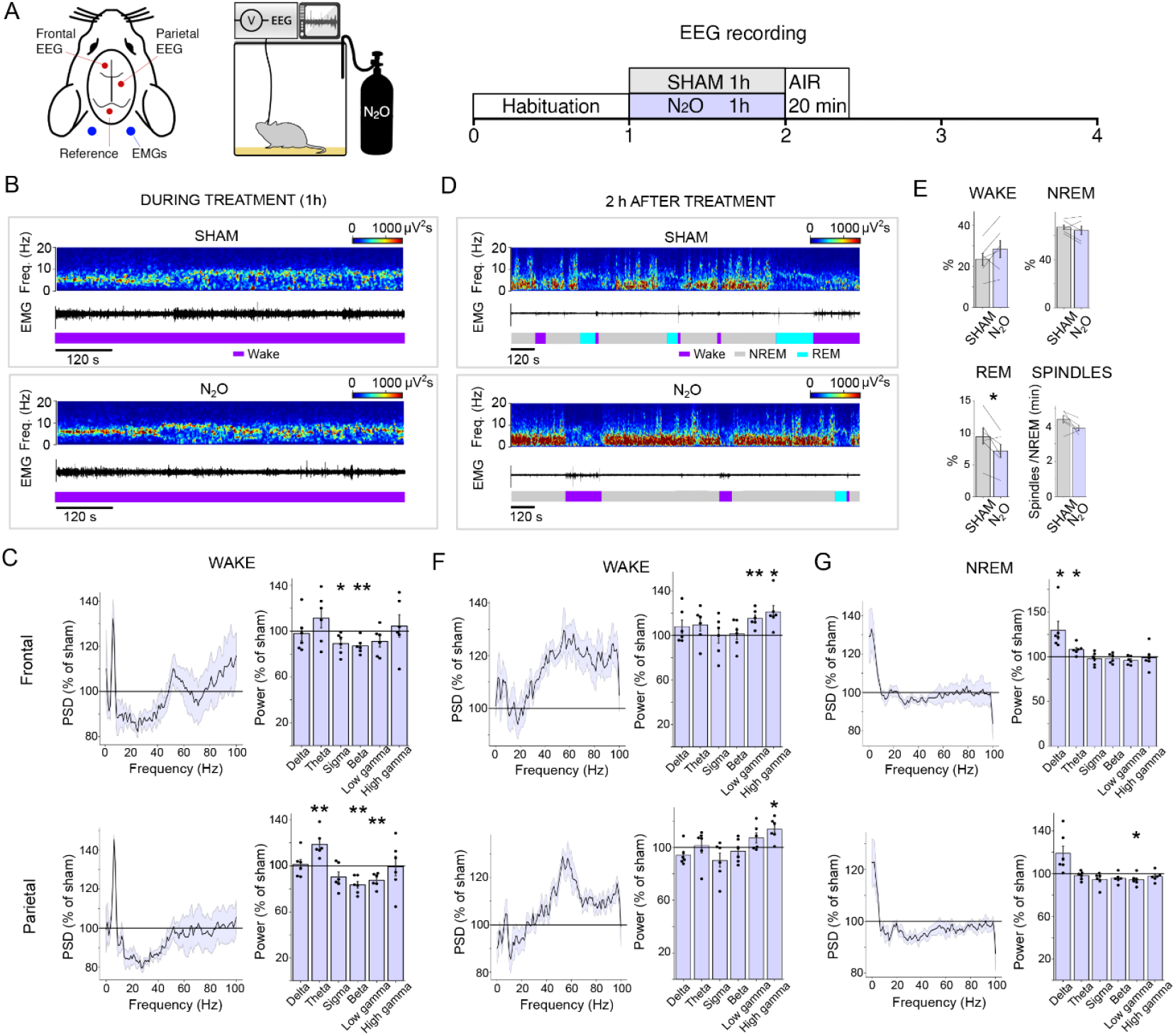
EEG recordings during and after 1h 50% N_2_O/O_2_ exposure. **A)** Schematic of the EEG recording screw electrode locations and timeline of the recording. Same mice were treated with N_2_O and sham with minimum of 1week washout between treatments. **B)** Example spectrogram demonstrating the frontal EEG derivation power spectral density during sham and N_2_O treatment, EMG activity and hypnogram. **C)** Power spectral density and power analyses from frontal and parietal derivations of wake EEG during N_2_O treatment as a percentage of the sham treatment values. Delta (0.5-4.5 Hz), theta (6-9Hz), sigma(10-15Hz), beta (15-30Hz), low gamma (30-50Hz), high gamma (50-100Hz). **D)** Example spectrogram demonstrating the frontal EEG derivation power spectral density after sham and N_2_O treatment, EMG activity and hypnogram. **E)** Percentage of wake, NREM and REM sleep, and the number of sleep spindles per minute of NREM sleep, during 2h after N_2_O and sham treatments. **F)** Power spectral density and power analyses from frontal and parietal EEG derivations of wake episodes 2h after N_2_O treatment as a percentage of the sham treatment. **G)** Power spectral density and power analyses from frontal and parietal EEG derivations of NREM sleep episodes 2h after N_2_O treatment as a percentage of the sham treatment. Abbreviations: EEG, electroencephalogram; EMG, electromyogram; Freq., frequency.; PSD, power spectral density. *:p<0.05, **:p<0.01.

We performed power spectral analysis when mice were awake during the treatments in both frontal and parietal EEG derivations (**Figure 3B-C**). During N_2_O treatment, wake theta (6-9Hz) power increased, while sigma (10-15Hz), beta (15-30Hz), and low gamma (30-50Hz) power decreased (**Figure 3C**) in relation to sham treatment. There was no significant difference in delta (0.5-4.5 Hz) or high gamma (50-100Hz) power. Next, we quantified the amount of wake, NREM and REM sleep as well as NREM sleep spindles during the 2h period following treatment (**Figure 3D-E**). REM sleep was slightly reduced, and there was a trend towards decreased sleep spindles following N_2_O treatment. The amount of wake or NREM sleep did not differ significantly from sham treatment. However, during wake episodes, both low and high gamma power were significantly increased following N_2_O treatment (**Figure 3F**). Moreover, NREM sleep delta power, also known as slow wave activity (SWA), indicative of sleep pressure and deep sleep, was significantly elevated after N_2_O treatment (**Figure 3G**).

These effects of N_2_O are reminiscent of subanesthetic ketamine, which has been shown to increase cortical gamma power following administration (Kohtala et al., 2019c, Kohtala et al., 2019a; Raith et al., 2020; Zanos et al., 2016) and to facilitate SWA during subsequent sleep in rodents (Feinberg and Campbell, 1995; Kohtala et al., 2019b; Koncz et al., 2023). Both increased gamma power and subsequent facilitation of sleep SWA have been hypothesized to reflect synaptic potentiation, which has been associated with the antidepressant effects of ketamine (Gilbert and Zarate, 2020; Kohtala et al., 2020; Rantamäki and Kohtala, 2020; Tononi and Cirelli, 2014).

### Nitrous oxide exerts antidepressant-like effects in the chronic corticosterone model

To evaluate the antidepressant-like properties of N_2_O, we employed a chronic corticosterone (CORT) stress model commonly used in preclinical antidepressant research. After several weeks of CORT exposure, mice received a single 1-hour treatment of 50% N_2_O/O_2_, mirroring the concentration and treatment time used in clinical trials (Nagele et al., 2021, Nagele et al., 2015). Behavior was then assessed in a battery of tests, including the open field test (OFT), the tail suspension test (TST), coat state score, the female urine sniffing test (FUST), sucrose preference test, as well as the 4-choice odor-based cognitive flexibility task (**Figure 4, Figure S3**).

**Figure 4:**
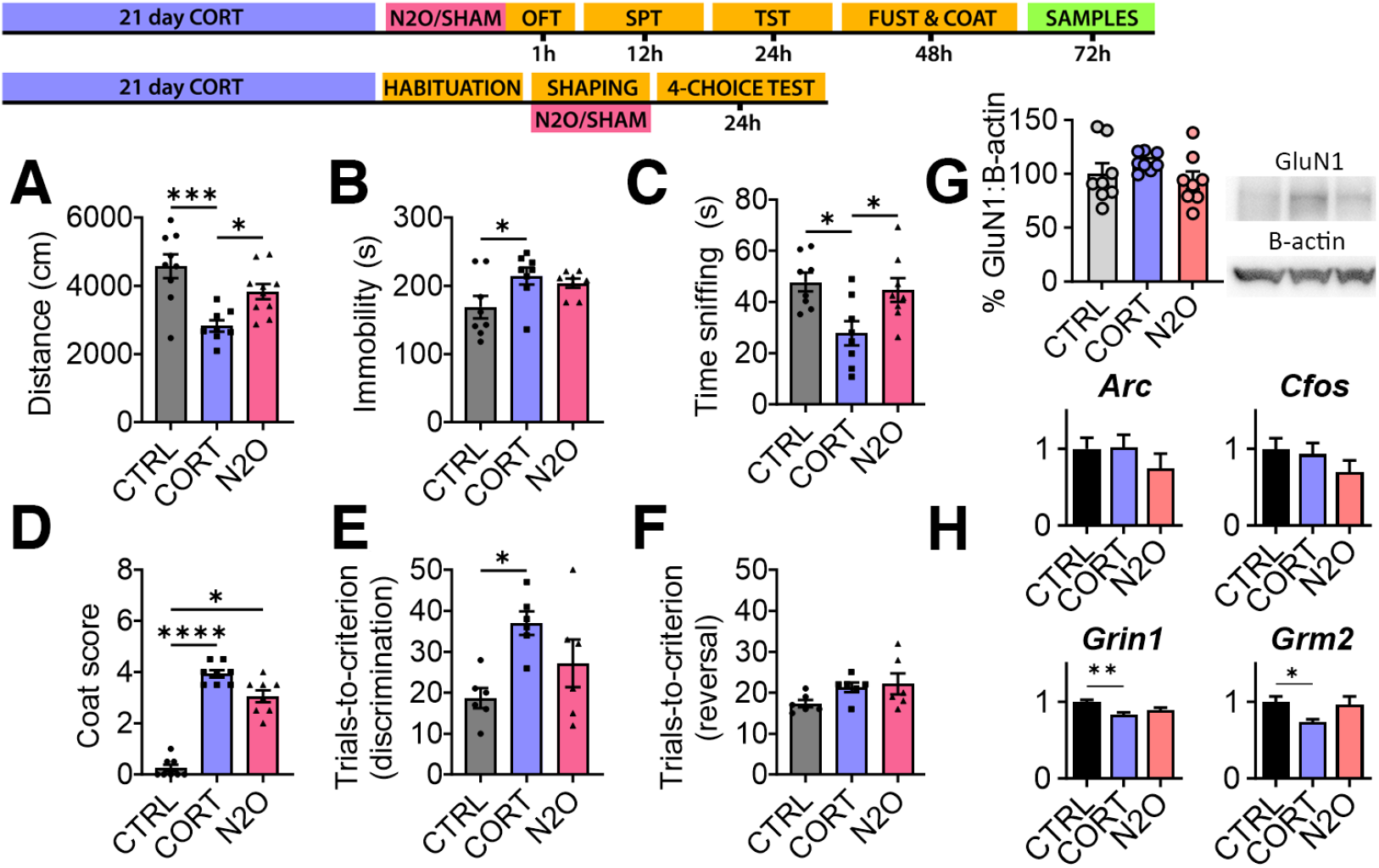
Behavioral and molecular effects of N_2_O treatment in the chronic corticosterone model. **A**) Total distance moved in open field test. **B**) Immobility time in tail suspension test. **C**) Time spent sniffing in the female urine sniffing test. **D**) Coat state score. **E**) 4-choice odor task discrimination trials. **F**) 4-choice odor task reversal trials. **G**) GluN1 protein levels in mPFC synaptosomes. **H**) mRNA expression in mPFC samples. *:p<0.05, **:p<0.01, ***:p<0.001, ****:p<0.0001

In the OFT, CORT-exposed mice receiving sham treatment exhibited a significant decrease in total distance traveled when compared to both naïve controls and N_2_O-treated mice, although center entries were reduced in both sham and N_2_O groups (**Figure 4A, Figure S3A**). In the TST, only sham-treated mice were significantly more immobile compared to controls (**Figure 4B**). In the FUST, sham-treated mice spent significantly less time sniffing female urine, consistent with anhedonia, while N_2_O treatment restored sniffing time to control-like levels (**Figure 4C**).

Coat state scoring, which reflects grooming and general well-being, was elevated in CORT-exposed mice prior to treatments, indicating a deteriorated coat condition consistent with the effects of stress exposure (**Figure S3B**). Although both sham and N_2_O groups remained higher than naïve controls 24 hours after treatment, there was a trend toward improvement in N_2_O-treated mice (**Figure 4D**). In contrast, only N_2_O-treated mice exhibited a significant drop in sucrose preference relative to controls (**Figure S3C**). Finally, in the 4-choice odor task, sham-treated mice required more trials to achieve criterion in the initial discrimination phase compared to controls, whereas N_2_O-treated mice were not significantly different from controls (**Figure 4E**). No differences emerged in the reversal phase (**Figure 4F**).

We also collected mPFC samples 72h after treatment and quantified the expression of key proteins and mRNA related to glutamatergic neurotransmission and synaptic plasticity (**Figure 4, Figure S4**). No statistically significant differences were found between the groups in the levels of synaptosomal GluN1 protein levels, which reflect synaptic NMDAR availability and have been reported to be modulated by corticosterone (Mikasova et al., 2017) (**Figure 4G)**. However, decreased mRNA expression of both *Grin1* and *Grm2*, which code for GluN1 and mGluR2, respectively, were observed in sham treated mice compared to controls. No significant differences were found in the expression of IEGs *Arc* and *Cfos* between the groups (**Figure 4H**). Moreover, no significant differences were found in the expression of *Gphn, Bdnf, Dlg4, Gria1, Grin2a, Grin2b, Grm5* (**Figure S4A**), or levels of BDNF protein in synaptic or cytosolic compartments (**Figure S4B**).

## Discussion

In this study, we investigated the molecular, cellular, and behavioral effects of a clinically used N_2_O concentration (50% N_2_O/O_2_) in mice. We focused on several key features that have been previously associated with the antidepressant effects of subanesthetic ketamine, such as cortical neuronal activation and wake–sleep EEG dynamics (Gilbert and Zarate, 2020; Kohtala et al., 2020). Our findings collectively indicate that N_2_O and ketamine share multiple mechanistic features, including rapid increases in cortical neuronal activity and modulation of wake and sleep EEG patterns. This convergence may provide insights into the common pathways underlying rapid antidepressant responses.

A key observation was that a brief acute exposure to 50% N_2_O/O_2_ robustly elevated calcium activity in the medial prefrontal cortex (mPFC). The calcium signals rose rapidly with N_2_O inhalation and subsided quickly upon cessation, mirroring the rapid pharmacokinetics of N_2_O (Quach et al., 2022). Because calcium influx in neurons largely occurs via NMDARs and voltage-gated calcium channels during action potentials (Chung, 2015), the observed calcium elevation most likely reflects a surge in neuronal firing. Subanesthetic ketamine has been reported to induce a rise in cortical calcium transients through disinhibition of excitatory networks (Ali et al., 2020; Hare et al., 2020; Kohtala, 2021; Moghaddam et al., 1997; Widman and McMahon, 2018), suggesting that a rapid increase in neuronal firing may constitute a shared mechanism for both ketamine and N_2_O. However, the synapsin-driven GCaMP6s expression used here does not distinguish between excitatory and inhibitory neuronal subpopulations, leaving open the question of whether cortical disinhibition also underpins the N_2_O response. Future studies targeting distinct neuronal types will be necessary to clarify this point.

Further supporting an N_2_O-induced enhancement of neuronal activity, we observed increased c-Fos protein and *Cfos* mRNA in the mPFC following N_2_O exposure. This upregulation of IEGs depends on a sustained calcium influx that initiates downstream signaling pathways, including MAPK, leading to transcriptional activation of *Cfos* and other activity-dependent genes (Chung, 2015). Our results align with prior findings of elevated c-Fos in several brain regions following N_2_O administration (Kaiyala et al., 2003; Kohtala et al., 2019c; Rozov et al., 2024) and corroborate the notion that N_2_O elicits a robust molecular imprint of neuronal excitation.

Along with molecular and cellular markers of cortical activation, we show that N_2_O modifies EEG activity both during and after treatment. Although we did not detect immediate increases in gamma power during the N_2_O exposure itself, we did observe elevated gamma activity and increased NREM slow-wave activity (SWA) during wake and sleep, respectively, in the 2-hour recovery period. These findings parallel reports of enhanced gamma power and subsequent augmentation of SWA following the administration of subanesthetic ketamine (Feinberg and Campbell, 1995; Gilbert and Zarate, 2020; Kohtala et al., 2020, 2019b, 2019a; Koncz et al., 2023). Notably, these effects have been hypothesized to reflect treatment-induced increases in synaptic potentiation (Duncan et al., 2013; Gilbert and Zarate, 2020; Rantamäki and Kohtala, 2020; Tononi and Cirelli, 2014). Interestingly, a recent study comparing subanesthetic ketamine and N_2_O also noted that only ketamine significantly increased acute gamma power, implying there may be divergent temporal profiles in how these agents modulate high-frequency oscillations (Rozov et al., 2024). Despite this difference, our results reinforce that N_2_O, like ketamine, produces longer-lasting alterations in cortical network dynamics that could be relevant for rapid antidepressant effects.

We extended our investigation to a chronic corticosterone (CORT) model of depression-like behavior to evaluate the potential antidepressant-like actions of N_2_O in mice. Consistent with emerging clinical findings (Nagele et al., 2021, Nagele et al., 2015; Rech et al., 2024; Shao et al., 2023; Yan et al., 2022), we found that a single 1-hour N_2_O treatment reversed some aspects of CORT-induced deficits. Notably, N_2_O improved open-field locomotion one-hour post-treatment and restored female urine sniffing to control levels 48 hours later, suggesting both rapid and sustained behavioral effects. Moreover, in the discrimination phase of the 4-choice odor task, only sham treated mice required significantly more trials to reach criterion when compared to controls, which could reflect effects of N_2_O on improving executive function as reported in depressed patients (Liu et al., 2023). However, differences were not observed in every behavioral measure, and some features of depression-like phenotype in CORT-exposed mice were not pronounced to begin with, which may have obscured potential treatment effects. Our results nonetheless support the notion that N_2_O exerts antidepressant-like effects in a stress-based animal model. Future work employing alternative models and behavioral assays may clarify the depth and duration of these antidepressant-like outcomes.

Finally, we examined mPFC tissue at 72 hours post-treatment to assess longer-term molecular changes linked to glutamatergic neurotransmission and synaptic plasticity. While we detected a significant reduction in *Grin1* and *Grm2* mRNA in CORT sham mice relative to controls, no notable alterations emerged in the levels of BDNF (protein or mRNA) or other synaptic proteins. The absence of robust alterations at the 72-hour time point is consistent with dynamic expression patterns that may normalize after initial upregulation. Indeed, prior work indicates that N_2_O can increase *Bdnf* expression at earlier time points (Kohtala et al., 2019b), suggesting that longer delays in analysis may fail to capture transient molecular shifts that accompany antidepressant responses.

## Conclusions

This study demonstrates that a single exposure to a clinically relevant concentration of N_2_O robustly increases neuronal calcium activity in the medial prefrontal cortex, which is accompanied by increased c-Fos expression. In vivo EEG measurements further reveal that N_2_O modulates cortical activity following treatment in both wake and sleep, specifically by enhancing gamma power in the waking state and delta activity during NREM sleep. These neurophysiological findings parallel those of subanesthetic ketamine, suggesting that shared mechanisms related to cortical activation and synaptic potentiation may underlie their rapid antidepressant effects. Consistent with previous clinical reports, a single N_2_O treatment also exerts antidepressant-like behavioral outcomes in a mouse chronic corticosterone stress model. Collectively, these findings highlight the utility of N_2_O as a valuable tool for probing the cellular and circuit-level processes underlying rapid antidepressant actions. Future investigations will be essential to delineate the specific neuronal populations and molecular pathways responsible for these effects.

## Acknowledgements

We are grateful to Tomi Rantamäki for support and valuable discussions. We thank Rui Rong Yang (Weill Cornell Medicine) for assistance with immunohistochemistry, Ryota Hasegawa (Weill Cornell Medicine) for advice on two-photon imaging, Jihye Kim (Weill Cornell Medicine) for advice on molecular and behavioral studies, and Jaakko Teppo (University of Helsinki) for providing access to computational resources.

## Author contributions

Conceptualization, S.K.; methodology, S.K., P.K.P., I.B., H.A.; investigation, S.K., P.K.P., I.B., H.A., L.I., P.K.; formal analysis, S.K., H.A., P.K, I.B..; funding acquisition, S.K, F.S.L., C.L.; project administration, S.K., F.S.L., C.L.; supervision, S.K., F.S.L., C.L.; writing – original draft, S.K.; writing – review & editing, S.K., P.K.P., H.A., F.L., C.L.

---

**Supplementary Figure 1.**
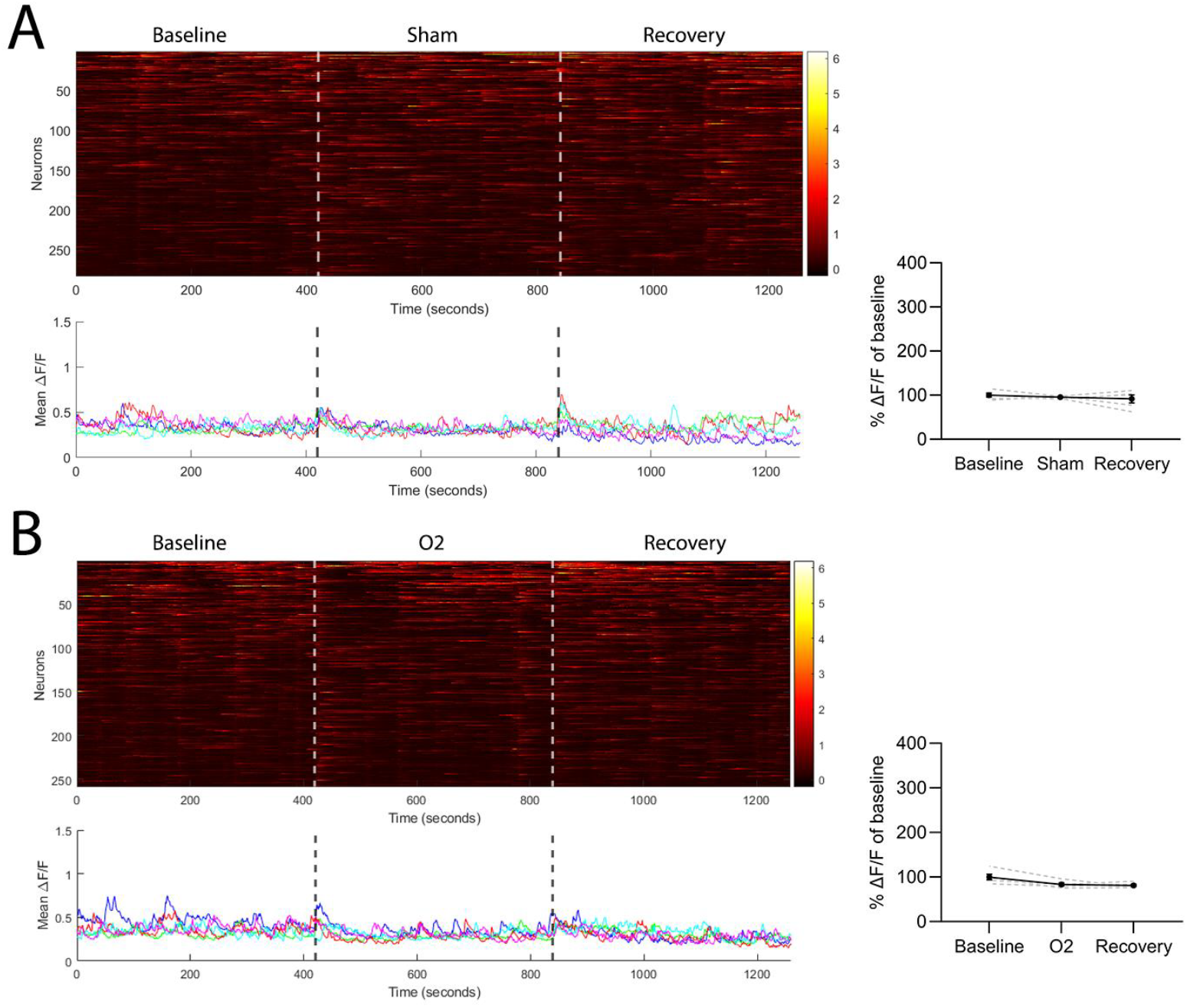
Neuronal mPFC calcium imaging during sham treatments. **A)** Representative heatmap of calcium traces throughout baseline (no gas), Sham (air flow), and Recovery (no gas) are displayed in the top figure. The plot below displays mean calcium signals from individual mice (n=5). On the right, plotted mean calcium activity corresponding to baseline, Sham, and Recovery periods across all subjects (n = 5), normalized to baseline. **B)** Representative heatmap of calcium traces throughout baseline (no gas), O2 (100% oxygen), and Recovery (no gas) are displayed in the top figure. The plot below displays mean calcium signals from individual mice (n=5). On the right, plotted mean calcium activity corresponding to baseline, O2, and recovery periods across all subjects (n = 5), normalized to the baseline period.

**Supplementary figure 2.**
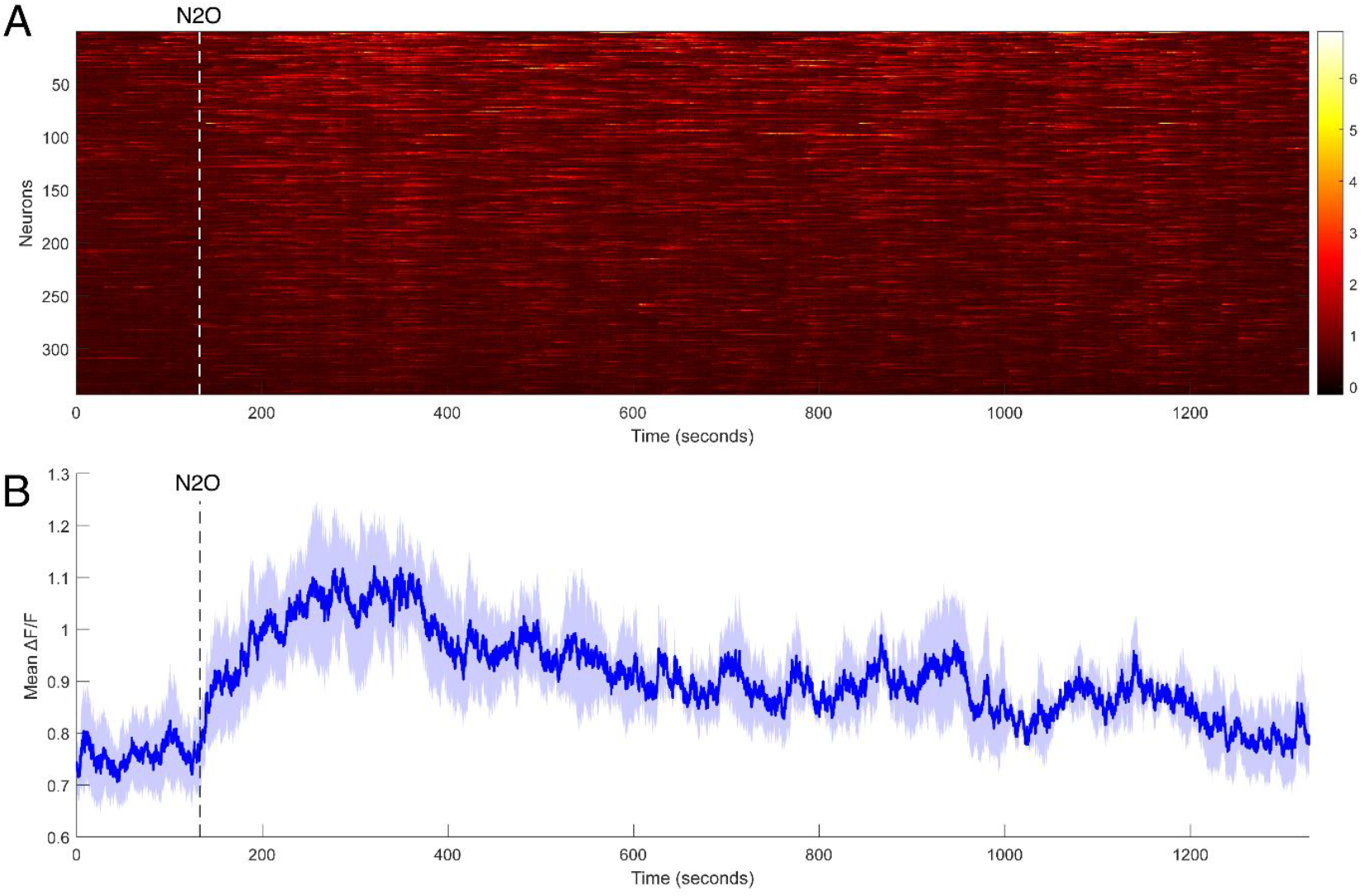
Neuronal mPFC calcium imaging during a continuous 50% N_2_O/O_2_ exposure. **A)** Representative heatmap depicting calcium activity in one subject across neurons. **B)** Mean calcium activity from 3 mice. The dark blue line depicts mean calcium signal and SEM is displayed as shaded light blue.

**Supplementary Figure 3:**
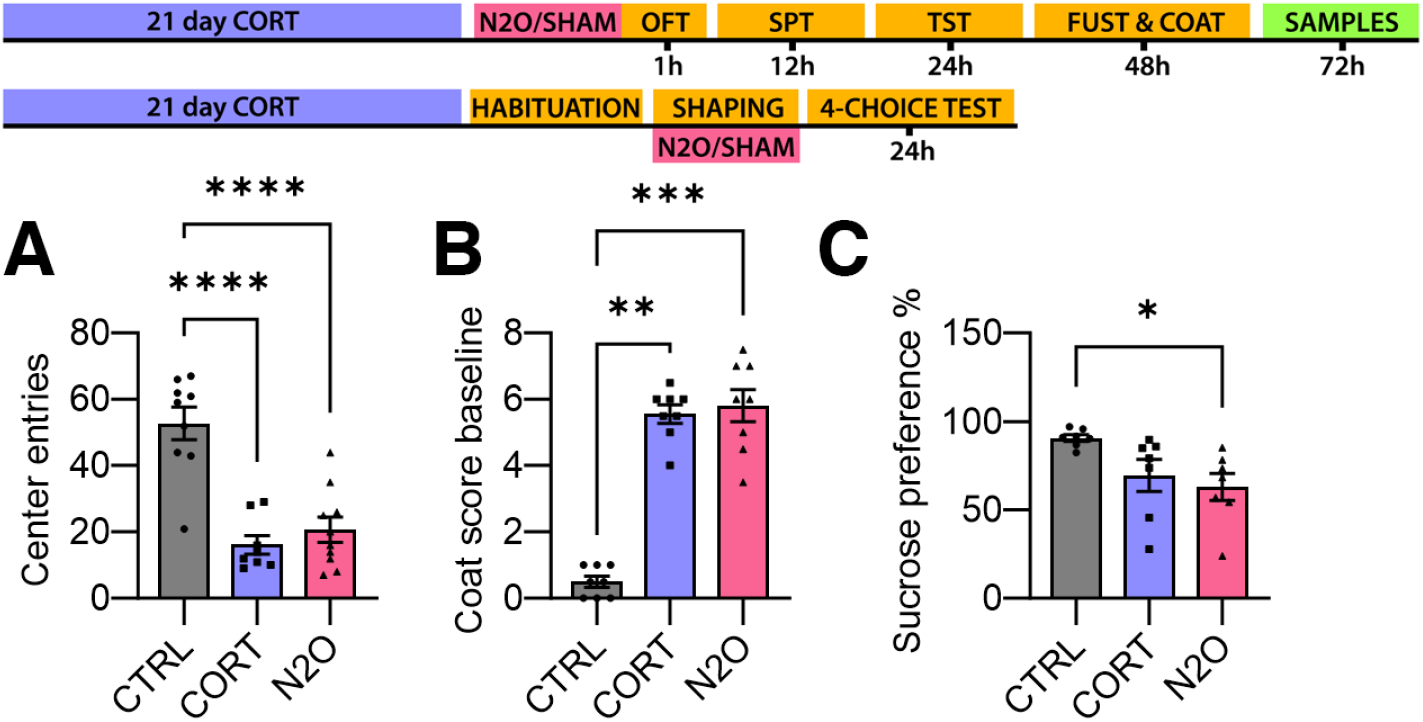
Behavioral effects of N_2_O treatment in the chronic corticosterone (CORT) model. **A)** Center entries in open field test. **B)** Coat score before treatment. **C)** Sucrose preference. *:p<0.05, **:p<0.01, ***:p<0.001, ****:p<0.0001

**Supplementary Figure 4:**
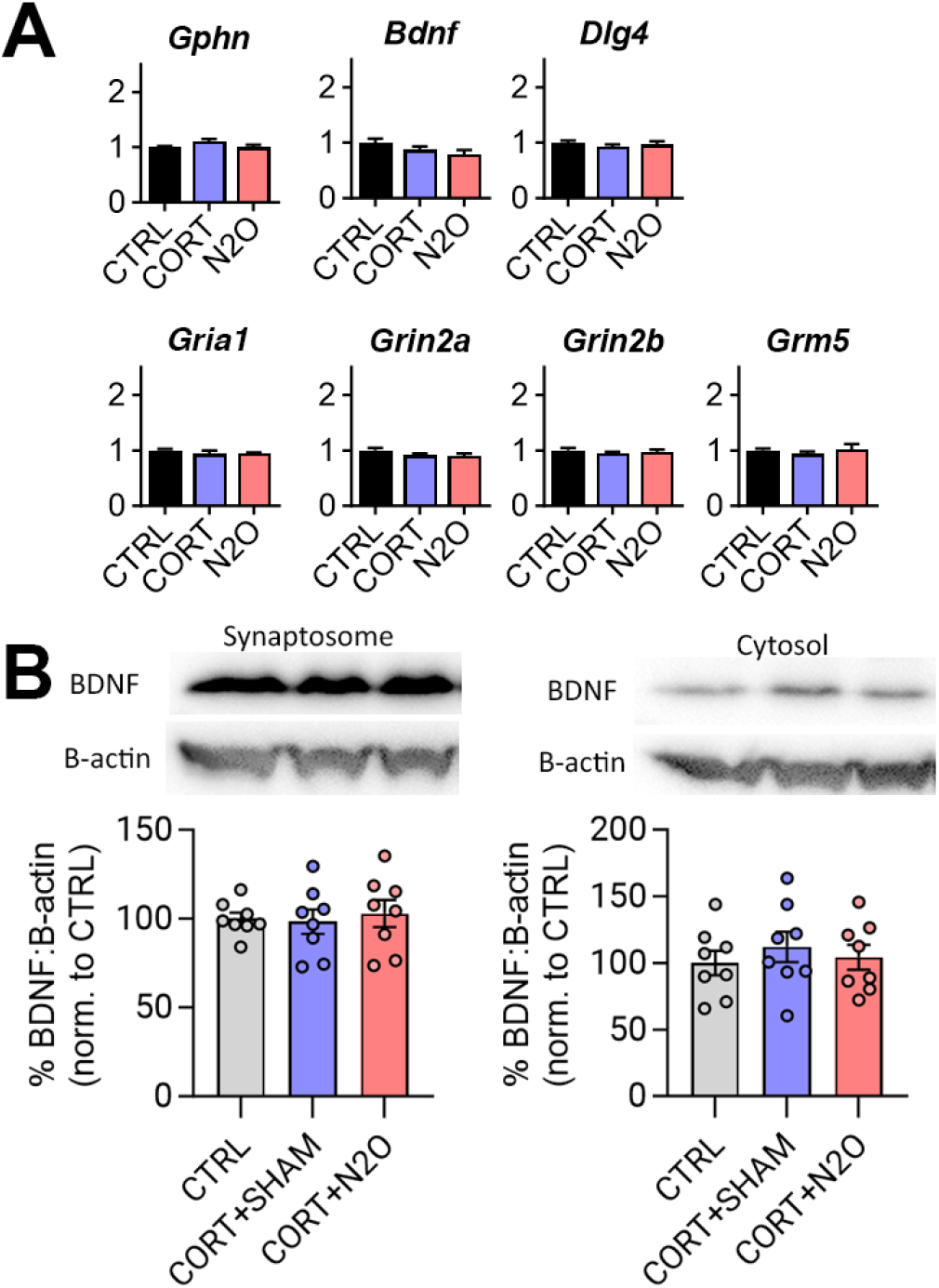
mRNA and protein changes in mPFC samples collected from chronic corticosterone (CORT) experiments 72 h after treatment. **A)** mRNA expression. Untreated controls (CTRL), sham treated CORT mice (CORT), N_2_O treated CORT mice (N2O). **B)** BDNF protein levels in synaptosomal and cytosolic extracts.

## Notes

### Competing Interest Statement

The authors have declared no competing interest.

